# The global exposure of species ranges and protected areas to forest management

**DOI:** 10.1101/2022.02.11.480150

**Authors:** Martin Jung, Matt Lewis, Myroslava Lesiv, Andy Arnell, Steffen Fritz, Piero Visconti

## Abstract

The majority of vertebrate species globally are dependent on forests, most of which require active protection to safeguard global biodiversity. Forests, however, are increasingly either being disturbed, planted or managed in the form of timber or food plantations. Because of a lack of spatial data, forest management has commonly been ignored in previous conservation assessments. Here we show – using a new global map of forest management - that disturbed and human managed forests cover the distributional ranges of most forest-associated species. Even more worrying, protected areas are increasingly being established in areas dominated by disturbed forests. Our results imply that species extinction risk and habitat assessments might have been overly optimistic with forest management practices being ignored. With forest restoration being in the centre of climate and conservation policies in this decade, we caution that policy makers should explicitly consider forest management.

## Introduction

Forests cover approximately 27% of the earth’s land surface (Buchhorn et al. 2020; Jung et al. 2020). They are the exclusive habitat of 54.5% of terrestrial vertebrate and many other plant, fungi and invertebrate species (Gibson et al. 2011; IUCN 2012; Hill et al. 2019), and can directly or indirectly benefit humankind through ecosystem services such as food or water, something particular relevant for the over 1.6 billion living within close proximity of a forest (Newton et al. 2020). Increases in human population and demand for food, non-timber and timber products, are resulting in forests in tropical, temperate and boreal regions being increasingly disturbed or modified by humans (Lewis et al. 2015; Curtis et al. 2018). Changes in forest use and management can affect the structural integrity of forests (Ghazoul et al. 2015; Lewis et al. 2015), ultimately reducing the size and connectivity of forest patches (Haddad et al. 2015) and affecting forest biodiversity (Hill et al. 2019). Yet, while a loss in forest cover can reduce local species richness (Melo et al. 2018) and increase the extinction risk of many species (Tracewski et al. 2016; Santini et al. 2019), it is not fully understood to what extent biodiversity is exposed to forest disturbances and management globally.

Forests are commonly disturbed and anthropogenically managed (Lewis et al. 2015). Forest disturbances can be caused by both natural causes (Thom & Seidl 2016), such as wildfires or insect outbreaks, and anthropogenic causes, such as selective logging and edge effects (Dantas de Paula et al. 2016; Matricardi et al. 2020), both of which can drive a forest to a ‘degraded’ state (Ghazoul et al. 2015; Chazdon et al. 2016). Edge effects include roads or nearby artificial land-use types that can reduce forest carbon biomass (Silva Junior et al. 2020) and affect local microclimates (Ewers & Banks-Leite 2013; Hardwick et al. 2015). Increasingly disturbed and degraded forests have become the focus of policy attention (Hansen et al. 2020; Newton et al. 2020), with a recent study having found that the amount of ongoing forest degradation already surpasses deforestation in the Brazilian Amazon (Matricardi et al. 2020). In addition to natural forest disturbance, many forests across the world are anthropogenically managed, for instance by active planting of forests for production of timber and non-timber products (Chazdon et al. 2016). Antropogenically exploited trees and timber plantations cover most of western Europe, Southern China, Japan and America (Jung et al. 2020), and agroforestry has long been recognized as a traditional form of land management, often using many native tree species (Zomer et al. 2016). Yet, the extent to which forest-associated biodiversity is exposed to different forest management types is unclear.

Owing to the reduction and simplification of structural complexity, disturbed and planted forests often have considerably lower biodiversity value (Chazdon et al. 2016). Disturbances and edge effects are commonly identified as a driver of worsening conditions in protected areas (Laurance et al. 2012), impacting local biodiversity (Pfeifer et al. 2017). And while (even exotic) forest plantations can potentially connect or form a tree-covered buffer around natural forest patches (Brockerhoff et al. 2008; Pellikka et al. 2009), there is mounting evidence that especially mono-culture plantations, such as pine or oil palm plantations, provide little or only reduced benefits for biodiversity (Farwig et al. 2008; Newbold et al. 2015). Although mixed, traditional management forms such as agroforestry can provide critical habitat (Hemp 2006; Bhagwat et al. 2008) and maintain a comparable high level of biodiversity (Jung et al. 2017), they also commonly have an altered species composition (Harvey & González Villalobos 2007). Yet, most current global forest pressure maps (Malhi et al. 2014; Lewis et al. 2015; Grantham et al. 2020) or frameworks for conservation or restoration assessments have ignored managed forests (Grantham et al. 2020; Hansen et al. 2020), or included them for a limited number of countries (Hill et al. 2019), presumably because of a lack of spatial data.

Remote sensing can assist in reliably identifying forest disturbances and management types. Fine-scale differences in remote sensing observations combined with visual evidence of selective logging or human structures nearby allow the separation of (visually) undisturbed from disturbed forests (Dantas de Paula et al. 2016; Curtis et al. 2018). Similarly, trees that were planted in regular spacing, such as timber or fruit plantations can be identified and delineated from high-resolution satellite imagery. Here previous studies have used single or multiple satellite observations to map the world’s intact forests (Potapov et al. 2008), small-scale disturbances caused by selective logging (DeVries et al. 2015) or regional gradients of different management (Pfeifer et al. 2016). Yet, until recently, no global remote-sensing derived maps of forest management types existed, with earlier attempts instead relying on several environmental predictors, little independent training or validation data (Schulze et al. 2019), or only being available at coarse scale (Curtis et al. 2018). The Nature Map Initiative has produced a new global high-resolution layer describing not only undisturbed and disturbed forests, but also several types of forest management identifiable from remote sensing.

In this study we investigate the exposure of forest-associated biodiversity to different types of forest management globally. Specifically, we combine estimates of the distribution of forest-associated vertebrate species with a novel, remote-sensing derived global map of forest management for the year 2015 (Fig. 1). We hypothesize that (*i*) the distributional range of forest-associated species is to a large degree covered by forests that are either disturbed or under some form of forest management, (*ii*) species threatened by extinction or threats associated with disturbances or forest extraction are disproportionately affected by parts of their range covered by disturbed or managed forests, and that (*iii*) protected areas are increasingly established in forests that cannot be considered undisturbed. Collectively, these hypotheses would suggest that several forest-associated species are confined to marginal intact habitats and addressing the management of these forests is critical to revert global biodiversity declines and improve the ecological state of forests globally.

**Fig. 1:**
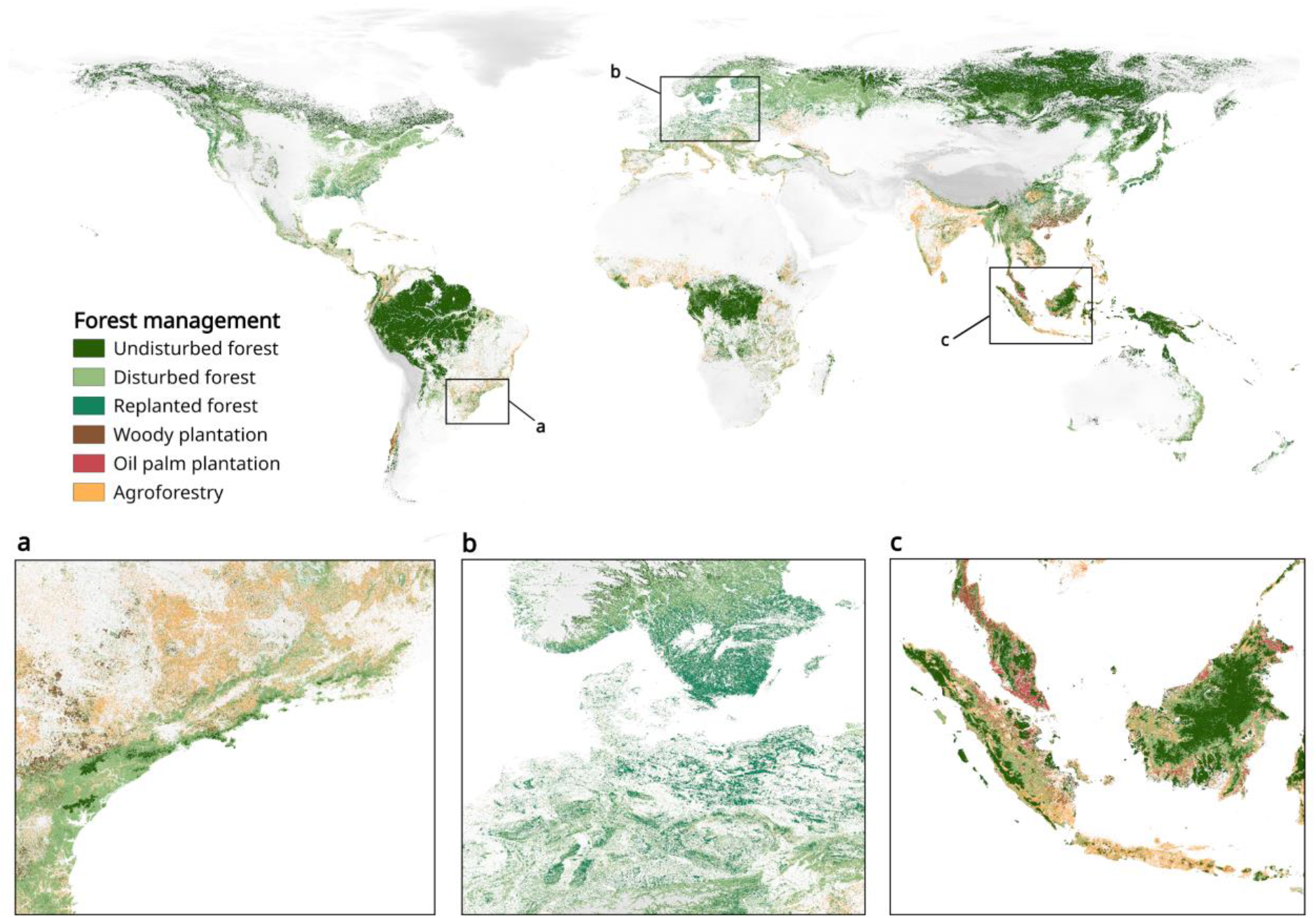
Global map of forest management types at ~100m resolution. Insets highlight the (a) remaining undisturbed forest in the Atlantic Forest region, (b) planted forests in central and northern Europe and (c) undisturbed forest amid palm oil and fruit plantations in Malaysia and Indonesia. Background shows a half-transparent Digital Elevation Model.

## Methods

Data on disturbed and planted forests came from a novel global forest management layer produced for the year 2015 at 100m resolution (Lesiv et al. submitted, 2020). The global forest management layer has in total six different classes, namely undisturbed (no visual signs of human impact), disturbed (visual impacts such as selective logging, clear cuts or built-up roads and human structures), and replanted forest (with a rotation period longer than 20 years), as well as woody plantations (with a rotation period of up to 15 years) and oil palm plantations, and agroforestry (which includes fruit tree plantations, shelterbelts or isolated trees on tropical pastures). We stress that the identification of managed forests was limited to those forms that are visually identifiable by remote sensing. The forest management layer was created entirely from remote sensing, combining high resolution training data, satellite time series and machine learning and shows overall good accuracy (81%) with independent validation data. The layer is described in full elsewhere and we refer to (Lesiv et al. submitted, 2020) for a more detailed description.

From the forest management layer we only considered plantations that had at least 10% tree cover fraction according to the global Copernicus Land cover product (Buchhorn et al. 2020) and following FAO definitions of forest. Opposed to other products of human impact on forests (Grantham et al. 2020), the forest management layer does not depend on any ‘scores’, stacking of arbitrarily selected land-use layers or definitions of ‘intactness’, but instead identifies forest management and disturbances directly from remote sensing. While this makes the mapped classes in our opinion more transparent, robust and replicable, we acknowledge that many forms of fine-scale forest disturbance can not reliably be detected from satellite imagery alone (Peres et al. 2006), which makes any estimates presented conservative.

For data on forest-associated vertebrate species distribution, we used spatial data on the ranges of amphibians (5,547), birds (8,434), reptiles (4,369, although we stress that not all reptiles globally have been assessed yet) and mammals (4,032) from the global IUCN Red List (ver 2019-2, (IUCN 2019)). We filtered the IUCN provided range data using standard criteria, e.g. by selecting only those parts of a species’ range where (i) it is extant or possibly extinct, 2) where it is native or reintroduced and 3) where the species is seasonally resident, breeding, non-breeding, migratory or where the seasonal occurrence is uncertain. Lastly, we limited our analyses only to those species that are ‘forest-associated’, which we define as any species for which ‘Forest’ is listed as known habitat preference according to IUCN. Lastly we obtained data on the threat status (e.g. CR, EN, VU, NT, LC, DD) of all selected species as well as - where available - data on IUCN listed threat types, such as for example ‘2.2 Wood & Pulp Plantations’ or ‘5.1 Hunting & trapping terrestrial animals’, which we broadly grouped into threat groups (See SI Table 1) and those with medium or high impact on a species.

In addition to data on the potential distribution of forest-associated vertebrate species, we also extracted similar statistics for all protected areas designated in or after 1995 available through the World Database on Protected Areas (IUCN & UNEP-WCMC 2020) from Google Earth Engine. We only selected established protected areas and furthermore excluded UNESCO-MAB Biosphere Reserves, following WDPA guidelines (Bingham et al. 2019).

We then summarized for each forest-associated species and protected area the amount of forest area (in ha) under each form of forest management. Protected areas which had no forest cover within their boundary were excluded from the analyses. To test whether forest area and management type differed among threatened (i.e. all CR, EN and VU) and non-threatened species, we used a logistic regression model fitted in a Bayesian framework using default uninformative priors (Bürkner 2018). Conditional model estimates were derived by summarizing the posterior in a mean estimate and 95% credible interval. We investigated model convergence by assessing the rhat statistic (all ~1.0) and the Markov chain Monte Carlo (MCMC) chains visually (SI Fig. 2). All data extractions and preprocessing were conducted on Google Earth Engine (Gorelick et al. 2017) and visualized in R (Wickham 2016; R Core Team 2019).

## Results

About 55% of the world’s forests were disturbed or managed in 2015. We found that 12,293 forest-associated vertebrate species (or 55.5% of all considered species) had disturbed or human managed forests as the most common type of forest within their range (Fig. 2, SI Table 2), and among reptiles, twice as many forest-associated species had most of their range now occupied by disturbed or planted forests (Fig. 2). Worryingly, forests within the ranges of 1,122 forest-associated species were predominantly of woody and oil-palm plantation and agroforestry type (SI Fig. 1, SI. Tab. 1).

**Fig. 2:**
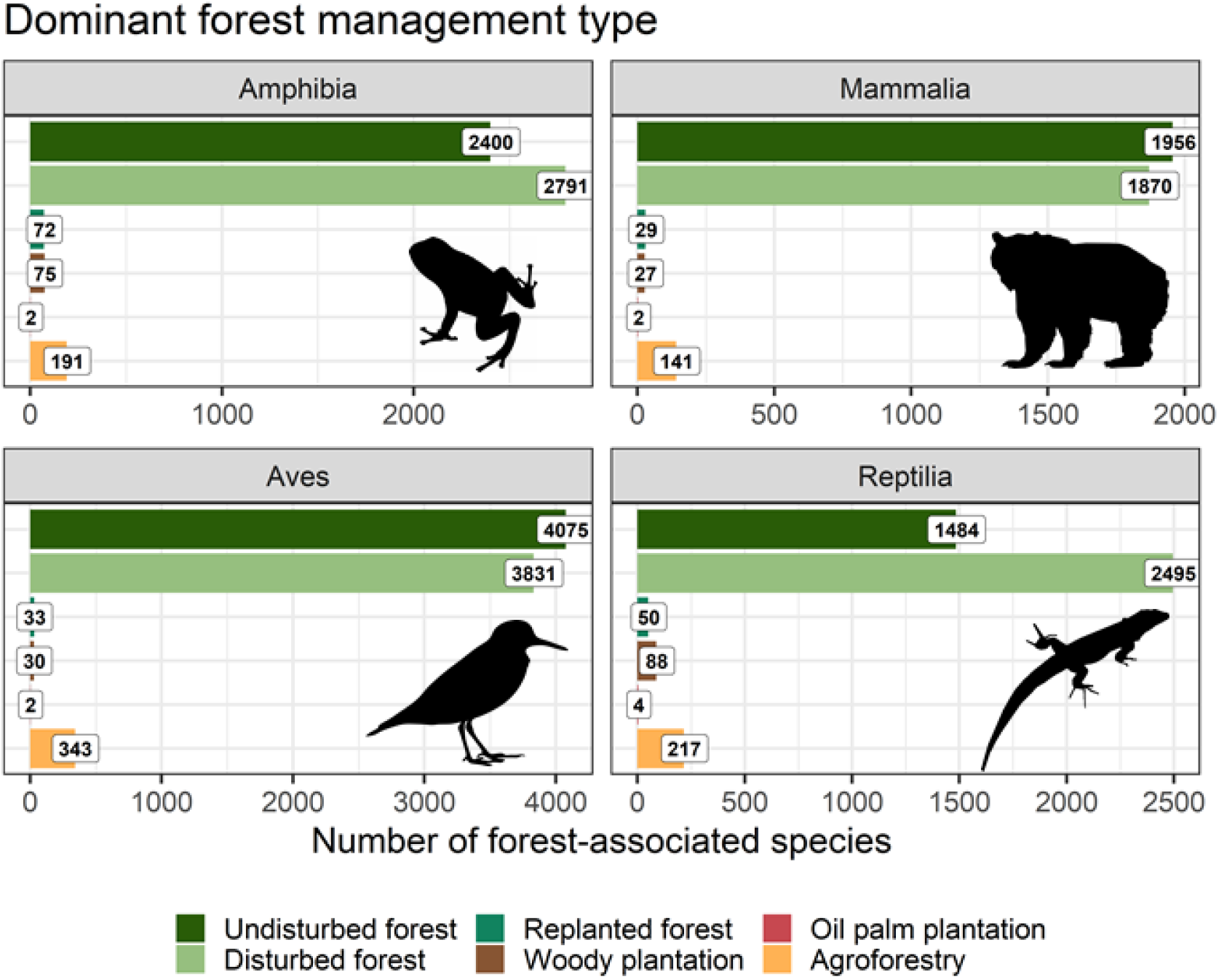
Dominant forest management type across all forested areas within each vertebrate species range. Numbered labels and x-axis show the total number of species. Colours and legend as in Fig. 1. Icons are public domain from phylopic.org

The amount of forest under different management types available to forest-associated species affected whether a species was classified as threatened by extinction. We found that an increase in forest area decreased extinction risk across all forest-associated species (SI Fig. 1). However species with a greater amount of undisturbed, disturbed and agroforestry forested areas in their range were more likely to be classified as non-threatened (Fig. 3a). In contrast, an increase in woody or oil palm plantation area did not decrease extinction risk probability nor did any difference in the amount of replanted forest (Fig. 3a). Species classified as non-threatened had overall larger amounts of undisturbed and disturbed forest within their range as well as a greater proportion of replanted small forest fragments present than for comparable threatened species (Fig. 3b, SI Fig. 3). Critically, the amount and distribution of forest area under different management types for data deficient species mirrored that of threatened species (Fig. 3b). If the distribution of unmanaged, disturbed and managed forests in a species range is any indication, this suggests that forest-associated data deficient vertebrate species are, in average, more likely to be at high risk of extinction than not.

**Fig. 3:**
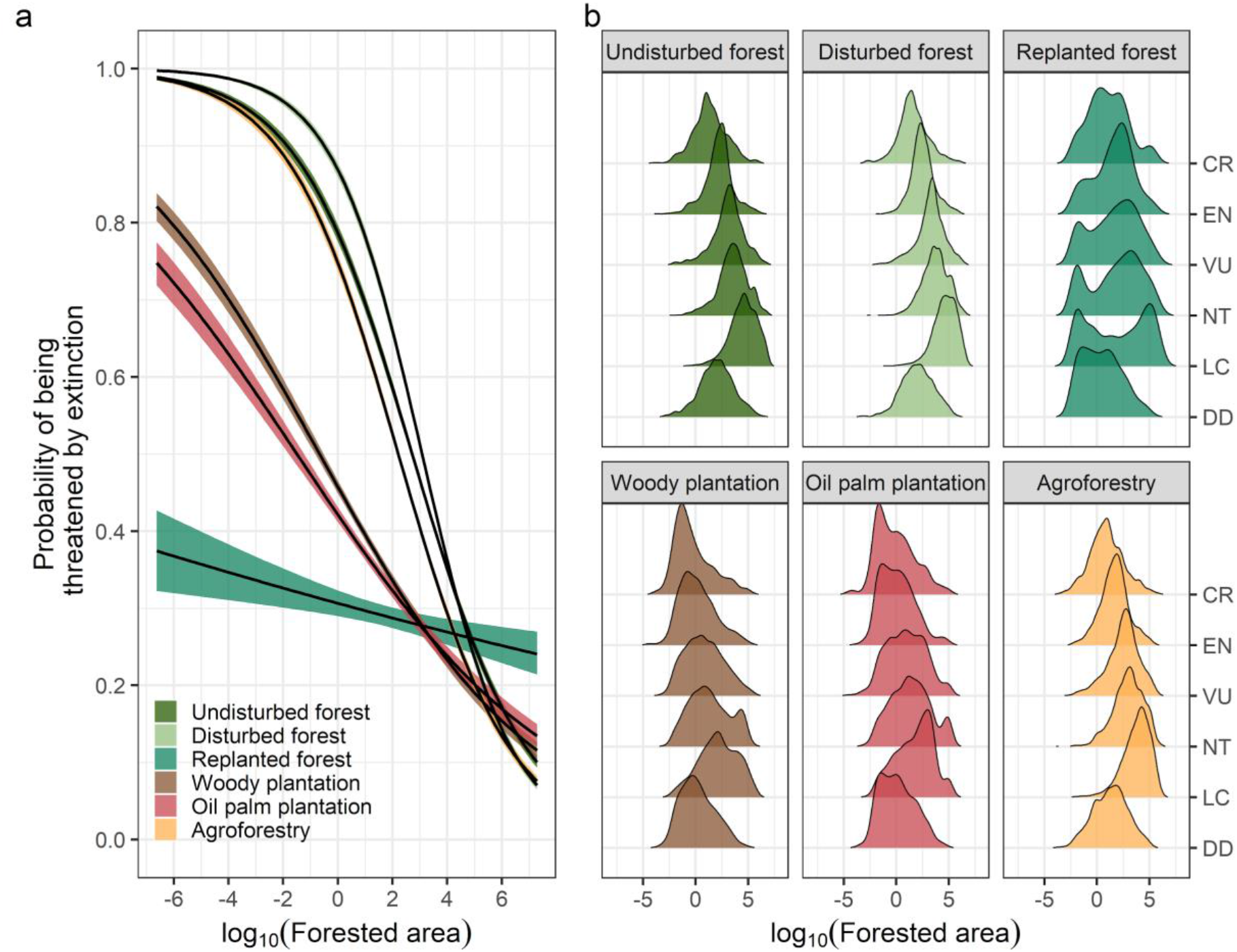
Marginal effect of an increase in forest area (log-transformed) on extinction risk probability, i.e. the probability that a species is classified as threatened according to IUCN. (**a**) Lines are mean estimates sampled from the model posterior with uncertainty bands showing the 95% credible interval. (**b**) Distribution of log10-transformed forested area estimates across species with different threat statuses according to IUCN. Colours as in Fig. 1

Furthermore, we found that, for species with available threat information, disturbed forests were the most common forest management type (SI Fig. 4). Agroforestry tended to be more often the dominant type of forest management within the range of species threatened by wood harvesting, persecution and subsistence farming (SI Fig. 4). Interestingly, many species which - according to IUCN - are strongly impacted by wood harvesting, did not have significantly more woody or fruit plantations in their ranges than the other forest management types.

Forests in terrestrial protected areas were under differing management types. Globally, protected areas contained 301 million ha of undisturbed forest (1.17% of all undisturbed forest), 121 million ha disturbed forest (0.5% of all disturbed forest) as well as 36.1 million ha of planted or managed forest (0.3% of all managed forest). Yet, irrespective of any IUCN assigned category of protection, the dominant forest management type within protected areas was disturbed forest, followed by replanted and then undisturbed forests (Fig. 4a). Interestingly, the majority of new protected areas designated between the years 2000 and 2010 are dominated by disturbed and replanted forest in the year 2015 (Fig. 4b), while few protected areas predominantly contain undisturbed forest. Predictably, few protected areas were established over predominantly woody or fruit plantations, indicating that protection measures mainly aimed at conserving forest that is not under intensive use by humans.

**Fig. 4:**
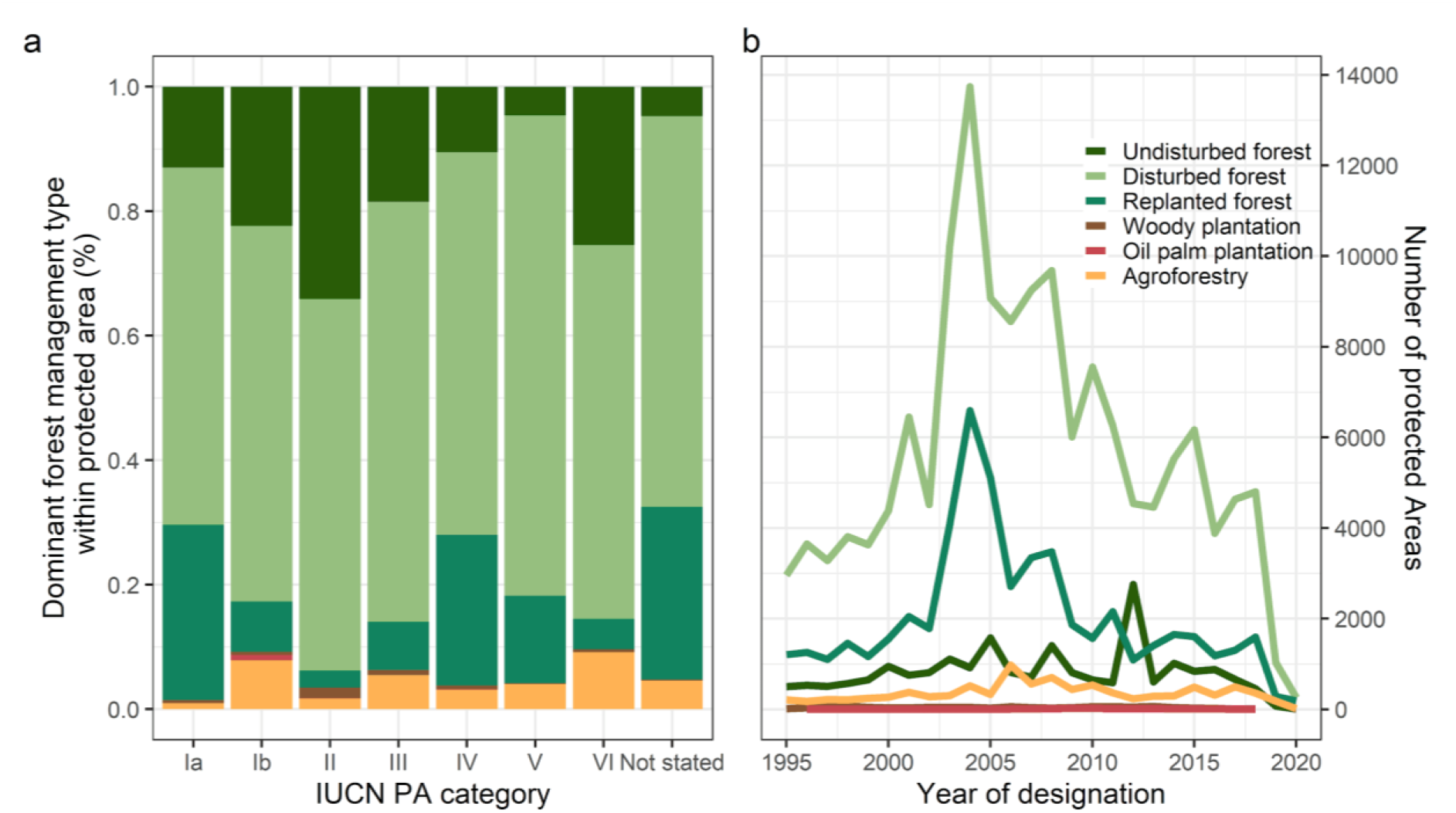
Dominant forest management type across (**a**) protected areas with different IUCN categorization and (**b**) number of newly designated protected areas in the last 25 years grouped by dominant forest management type. Colours as in Fig. 1.

## Discussion

Humans have altered the majority of forests across the world, with 55% of forests being either disturbed or managed by humans. Our results show that over half of the ranges of forest-associated vertebrate species across the world are covered by either disturbed or human managed forests (Fig. 2), with the amount being particularly high for species threatened by extinction (Fig. 3). Furthermore, we show that many designated protected areas are already dominated by disturbed and replanted forests (Fig. 4), highlighting both the value of past forest restoration measures as well as the need to step up protection of remaining undisturbed forests.

Replanting forest is considered to be a primary target for restoring degraded habitats. Interestingly, our results indicate that increasing or decreasing the amount of planted forest within forest-associated species ranges has little influence on whether the species is currently classified as threatened by extinction (Fig. 3). This could indicate that most previous forest restoration efforts have either not yet explicitly benefitted forest-associated vertebrate species, or lag effects due to outdated IUCN assessments or past land use change affect the conservation status (Chazdon et al. 2008; Jung et al. 2019; Veldman et al. 2019). For example, areas previously covered by native tree species in Kenya have been increasingly afforested using exotic pine trees, often with little benefit for native species (Farwig et al. 2008; Pellikka et al. 2009). Human planted forests are not necessarily bad for biodiversity (Carnus et al. 2006), they are in fact essential if we are to subject large tracts of degraded, previously forested land to habitat restoration (Chazdon 2008; Chazdon et al. 2008) and climate mitigation efforts. Yet those planted forests need to be established in places where they do not displace natural habitats, such as forests or savannas (Veldman et al. 2019), or native tree species, and do not negatively impact the livelihood of local communities in developing countries (Malkamäki et al. 2018). Thus, further afforestation and reforestation efforts should be carefully evaluated with regards to local contexts and their potential benefits for biodiversity conservation.

Our results also have important implications for conservation applications that use species habitat preferences and land-cover maps to refine species ranges to Area of Habitat (AOH) maps (Brooks et al. 2019). Because, most existing AOH use exclusively land cover products (Rondinini et al. 2011; Ficetola et al. 2015), thus ignoring forest management, it follows that AOH might be grossly overestimated if populations of forest-associated species are not able to persist in disturbed or managed forests. Novel hybrid maps have been developed that alleviate some of these issues by accounting for both land-cover and land-use (Jung et al. 2020), however, these maps do not thematically consider all possible forms of management that might be relevant for ecological or conservation studies. We suggest that more evidence is needed on the persistence of forest-associated species in disturbed and managed forests to ensure that maps of habitat-based refinements are fit for purpose.

While the global forest management map is the most detailed spatial-explicit quantification to date, we acknowledge that not all forms of anthropogenic disturbances can likely be detected from remote sensing (Peres et al. 2006), thus our estimates will likely be an underestimate. This is exemplified by the fact that although many forest-associated species are known to be sensitive to anthropogenic threats (Maxwell et al. 2016), we found few differences between species threatened by disturbances or wood harvesting (SI Fig. 4). We can also not rule out that some types of forests have been misclassified, which can impact our analyses (Sexton et al. 2016; Estes et al. 2018). Furthermore, we also highlight that our analysis does not take into account species occurrence and relative abundance across forest management types (we performed only range overlaps) and many - particularly disturbance sensitive - species do not necessarily inhabit all forests everywhere (Pfeifer et al. 2017). More work is needed on the impact of disturbances and wood harvesting on species local occurrence, population density and persistence, as well as more detailed mapping of forest management types at national and regional scales.

As we move into a decade of ecosystem restoration, we urge conservationists and policy makers to consider different types of forest management. Critically, ignoring forest management and focussing on forest cover alone, can give the misleading impression of no-net forest loss when in fact native, undisturbed forests are being replaced by woody plantations or getting disturbed (Tropek et al. 2014). With an increasing proportion of the Earth’s forests being disturbed or managed, we need to better account for and investigate the impact of forest management on the persistence of species populations and the effectiveness of conservation efforts.

## Code availability

Code used for the analysis and extracted data will be made openly available upon acceptance <To be inserted >

## Data availability

The global forest management layer will be made openly available as part of another article. Data on the distribution of vertebrate species and protected areas can be requested from the respective data providers, namely IUCN and Birdlife International. Data on threats status and existing threats are available from the IUCN Red List. Extracted data for each species is made available in SI Table 2 and the code repository.

