## Supplementary Information for "The global exposure of species ranges and protected areas to forest management"

**Running Title:** Conservation and forest management

Authors: Martin Jung1*, Matt Lewis1, Myroslava Lesiv2, Andy Arnell3, Steffen Fritz2 & Piero Visconti1

1 Biodiversity Ecology and Conservation Research Group, International Institute for Applied Systems Analysis (IIASA), Schlossplatz 1, A-2361 Laxenburg, Austria
2 Novel Data Ecosystems For Sustainability Research Group, International Institute for Applied Systems Analysis (IIASA), Schlossplatz 1, A-2361 Laxenburg, Austria
3 UN Environment Programme World Conservation Monitoring Centre (UNEP-WCMC), 219 Huntingdon Road, Cambridge CB3 0DL, United Kingdom

### Supplementary Information:

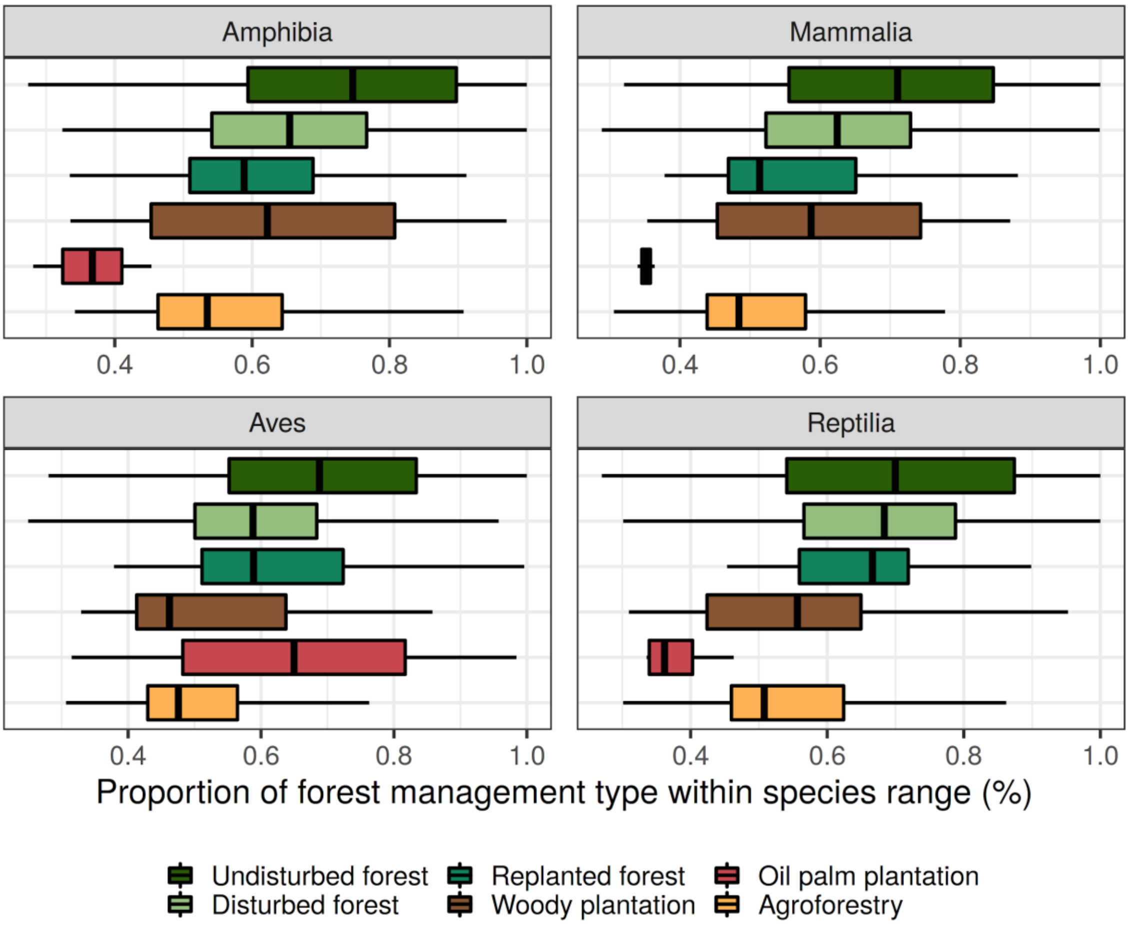

SI Figure 1: Distribution of forest management type estimates within a species range when non-occurring types are ignored. Values show the proportion of each class with boxplots indicating the lower, median and upper quantiles. Colours and legend as in Fig. 1.

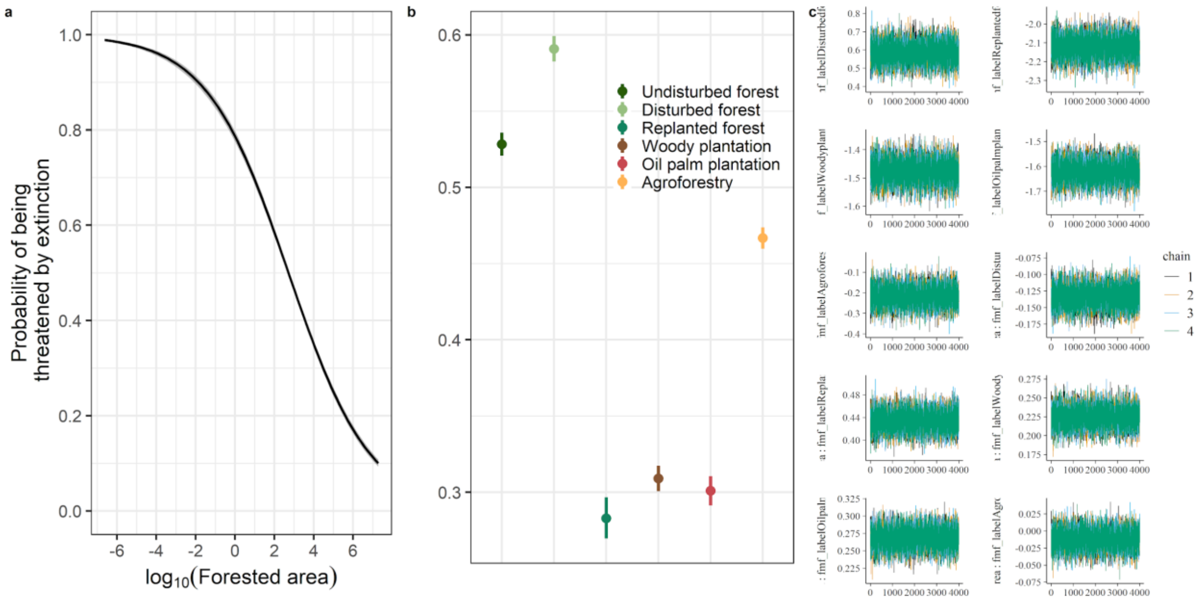


SI Figure 2: Posterior marginal effects of the logistic model (see methods) individually for (**a**) log-transformed forest areas and (**b**) mean effect and credible intervals of forest management on extinction risk. (**c**) Traceplot of Markov chain Monte Carlo draws across all four fitted chains shown for all fixed effects (minus the intercept that is undisturbed forest) and the fixed effects in interaction with log-area.

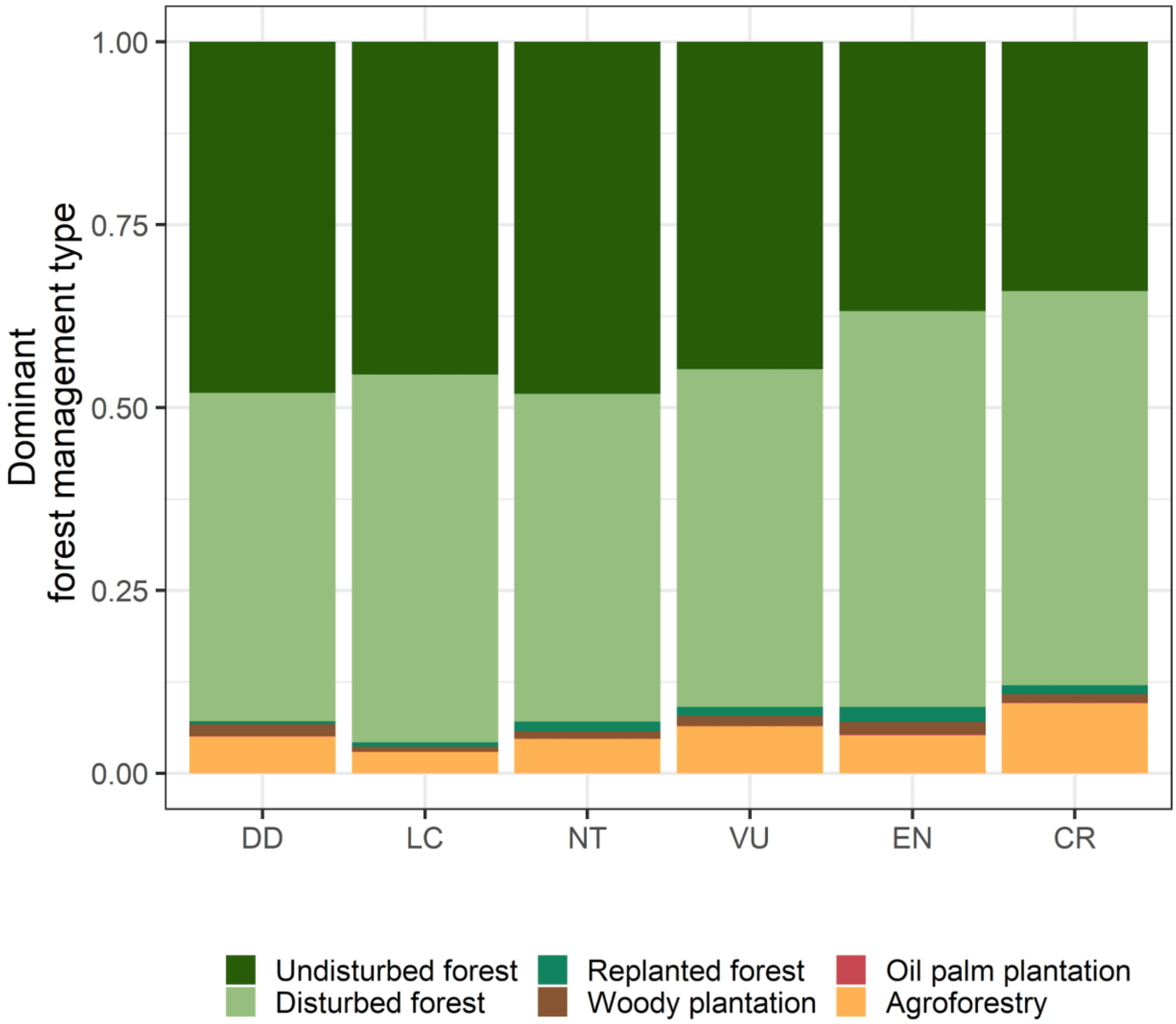


SI Figure 3: Proportion of each forest management type that is dominant within a species range. Colours as in Fig. 1

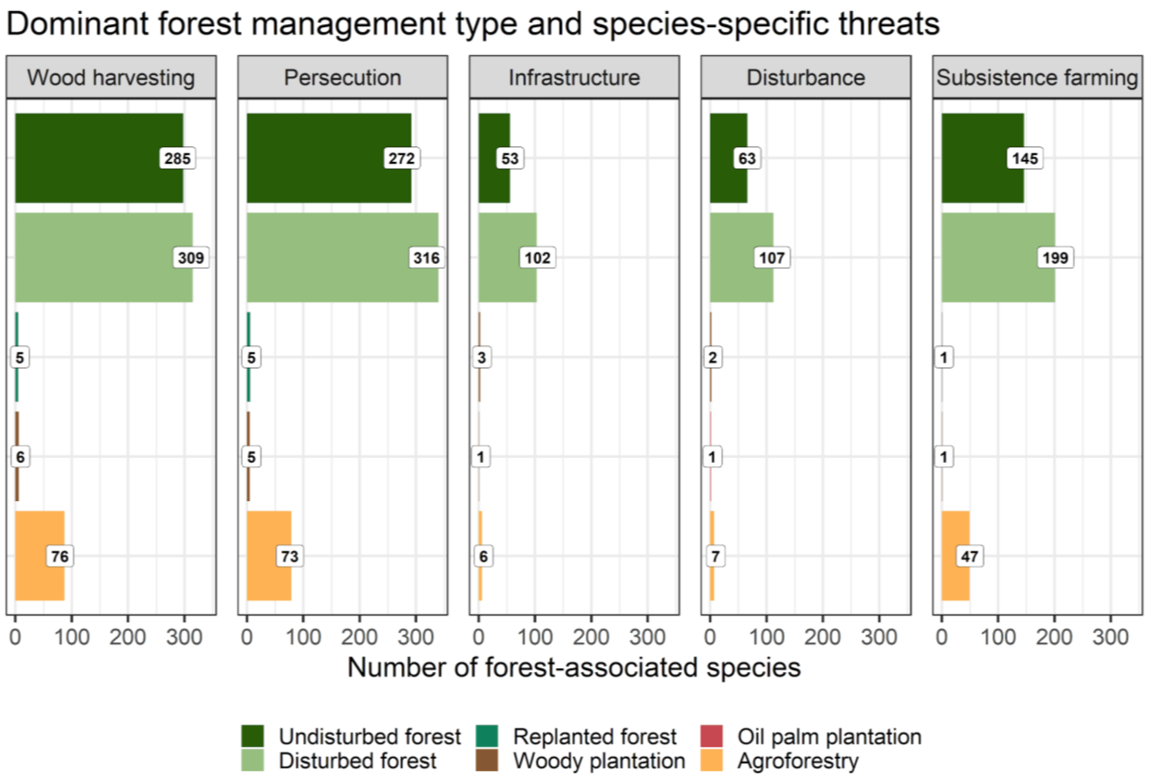


SI Figure 4: Number of forest-associated species in relation to species-specific threats with reported medium or higher impact on the species (see methods). Grouped by forest management type and coloured as in Fig. 1.

SI Table 1: Matching table between IUCN threat categories and the groups used in the analysis.

| **Threat_grouping** | **IUCN_threat** |
|---|---|
| Disturbance | Motivation Unknown/Unrecorded |
| Disturbance | Unintentional effects (species is not the target) |
| Disturbance | Work & other activities |
| Disturbance | Noise pollution |
| Disturbance | Tourism & recreation areas |
| Disturbance | Motivation Unknown/Unrecorded (Batrachochytrium dendrobatidis) |
| Disturbance | Light pollution |
| Disturbance | Recreational activities |
| Infrastructure | Housing & urban areas |
| Infrastructure | Roads & railroads |
| Infrastructure | Roads & railroads (Vulpes vulpes) |
| Infrastructure | Housing & urban areas (Batrachochytrium dendrobatidis) |
| Persecution | Gathering terrestrial plants |
| Persecution | Unintentional effects: (subsistence/small scale) [harvest] |
| Persecution | Intentional use: (subsistence/small scale) [harvest] |
| Persecution | Intentional use (species is the target) |
| Persecution | Hunting & trapping terrestrial animals |

| | |
|---|---|
| Wood harvesting | Logging & wood harvesting |
| Wood harvesting | Agricultural & forestry effluents |
| Wood harvesting | Unintentional effects: (large scale) [harvest] |
| Wood harvesting | Intentional use: (large scale) [harvest] |
| Wood harvesting | Logging & wood harvesting (Batrachochytrium dendrobatidis) |
| Wood harvesting | Wood & pulp plantations (Batrachochytrium dendrobatidis) |
| Wood harvesting | Wood & pulp plantations |
| Subsistence farming | Small-holder farming |
| Subsistence farming | Small-holder plantations |
| Subsistence farming | Nomadic grazing |
| Subsistence farming | Small-holder grazing, ranching or farming (Batrachochytrium dendrobatidis) |
| Subsistence farming | Small-holder grazing, ranching or farming (Batrachochytrium salamandrivorans) |
| Subsistence farming | Small-holder farming (Batrachochytrium dendrobatidis) |
| Subsistence farming | Shifting agriculture (Batrachochytrium salamandrivorans) |
| Subsistence farming | Small-holder farming (Batrachochytrium salamandrivorans) |
| Other | Other |

SI Table 2: Dominant proportion of each forest management type within the range of forest-associated species according to IUCN. Columns give the IUCN id number (id_no), forest management type and proportion with regards to the total forest area within the range.

<Supplied separately>
